# Clinical pattern of antibiotic overuse and misuse in primary healthcare hospitals in the southwest of China

**DOI:** 10.1101/585257

**Authors:** Chang Yue, Sarunyou Chusri, Rassamee Sangthong, Edward McNeil, Hu Jiaqi, Du Wei, Li Duan, Fan Xingying, Zhou Hanni, Virasakdi Chongsuvivatwong, Tang Lei

## Abstract

**Purpose:** Overuse and misuse of antibiotics are the primary risk factors for antibiotics resistance. Inadequate professional competence of primary care physicians might exacerbate these problems in China. This retrospective study aims to document the clinical pattern of antibiotics use and its overuse and misuse rates in rural primary care institutions, and to evaluate the association between antibiotics use and characteristics of the physicians and their patients.

**Methods:** Medical records from 16 primary care hospitals in rural areas of Guizhou province, China were obtained from the Health Information System in 2018. Classification of incorrect and/or unnecessary use, escalated use and combined antibiotics use was based on the Guiding Principle of Clinical Use of Antibiotics (2015, China) and the standard of USA Centers for Disease Control and Prevention. Generalized Estimating Equations were employed to determine predictive factors for inappropriate antibiotics use.

**Results:** A total of 74,648 antibiotics prescriptions were retrieved. Uncomplicated respiratory infection was the most common disease accounting for 58.6% of all prescriptions. The main antibiotic group used was penicillins (51.5%) followed by cephalosporins and macrolides (14% each). Of 57,009 patient visits, only 8.7% of the antibiotic prescriptions were appropriate. Combined, escalated, and incorrect and/or unnecessary antibiotics use was found in 7.8%, 6.2% and 77.3% of patient visits, respectively, of which 28.7% were given intravenously. Antibiotics misuse was significantly more likely among newly employed physicians with lower levels of professional education. Adult patients and those who had public insurance had a higher risk of being prescribed incorrect and/or unnecessary antibiotics.

**Conclusion:** Overuse of antibiotics for uncomplicated respiratory infection and use of cephalosporins, macrolides and injection antibiotics in primary care are the major problems of clinical practice in rural areas of Guizhou.

## Introduction

Antibiotics resistance is a growing global public health issue [1]. Between 2000 and 2010 global antibiotics consumption grew by more than 50% based on data from 71 countries, including China [2]. Excessive consumption of antibiotics mainly leads to the development of antibiotics resistance. China consumes the second largest amount of antibiotics in the world [3, 4] with a prescription rate twice that recommended by the World Health Organization (WHO) [5] and rural areas have higher rates than urban areas [6]. In this regard, the Chinese government has introduced a number of regulations in the last decade, but it has not played a significant role in rural areas [7].

The majority of people in southwest China live in rural areas where primary care physicians usually provide the health services [8]. Most physicians there have non-degree training yet are allowed to prescribe antibiotics in the national list due to personnel shortages. Previous studies reported a high irrational antibiotics prescription use among primary care physicians [7, 9]. Strengthening the knowledge and practice of rational use of antibiotics among primary care physicians is one way to reduce antibiotic resistance. Thus, we need to understand how antibiotics are unnecessarily used, for example prescribed with incorrect choices for particular diseases, escalated use (prescribing more expensive and broad spectrum antibiotics when cheaper and more specific antibiotics can give the same result) and unnecessary use of antibiotics given intravenously.

The aim of this study was to document the clinical patterns of antibiotics prescriptions in a rural primary care setting where physicians are mostly non-degree trained. The secondary objective was to detemine the association between antibiotics use and various characteristics of patients and physicians.

## Materials and methods

Ethical approval: This research was approved by the Institutional Review Board of Prince of Songkhla University, Thailand (REC: 60-285-18-5). Ethics committees approved the consent procedure. All participants provided their written informed consent to participate in this study.

### Study setting

Out of 1,399 township public hospitals in Guizhou rural areas, 132 use the same health information system (HIS) developed by the Department of Public Health. Those that had more than three outpatient physicians were randomly selected to participate in the study.

### Data retrieval process

Only data from outpatient units were retrieved. Demographic characteristics of the patients, and education and work experiences of the physicians were obtained from the Personnel Management Department. All patients prescribed antibiotics during February to August, 2018 were included in the analysis. The primary diagnoses of all diseases except for tuberculosis (since the treatment options are fixed and standardized), were grouped into 10 diagnostic categories according to the international classification for diseases version 10 (ICD-10) code [10]. Antibiotics were classified, based on the 2018 National Catalogue of Clinical Application of Antibacterial Drugs, into seven groups: penicillins, cephalosporins, macrolides, quinolones, lincosamides, nitroimidazoles, and aminoglycosides [11]. We focused on systemic antimicrobials excluding topical antimicrobial prescriptions such as ophthalmic ointments and skin creams.

Based on the US Centers for Disease Control and Prevention (CDC) [12], appropriate use of antibiotics refers to prescribing the right group of antibiotics for the given diagnosis, at the right dose, at the right time, and for the right duration. However, in this study, we defined appropriate use according to the Guiding Principle of Clinical Use of Antibiotics (2015, China) and the standard of the US CDC [13, 14]. Inappropriate antibiotic use was categorized into 1) incorrect and/or unnecessary use, defined as prescribing antibiotics without a clear indication, 2) escalated use, defined as using a broad-spectrum instead of a narrow-spectrum antibiotic for any given disease, and 3) combined use, defined as use of more than one antibiotic group per patient visit. In the study, the term “misuse” of antibiotics was used to describe incorrect and/or unnecessary use and the term “overuse” used to describe escalated or combined use.

### Data analysis

The units of analysis were both antibiotic prescriptions and patient visits where one physician may prescribe several antibiotics to a patient on any given day but a patient can have only visit one doctor per day. Cross-tabulation between the groups of antibiotics and the disease categories by ICD-10 diagnosis code was used to determine the pattern and appropriateness of antibiotics use. Misuse of antibiotic prescriptions was aggregated by antibiotic group. Univariate odds ratios were calculated to determine factors associated with inappropriate antibiotic use. To account for the correlation of antibiotic prescriptions by the same physician, multivariate modelling using Generalized Estimating Equations (GEE) was employed to determine factors associated with inappropriate antibiotic use. R version 3.3.1 was used for all data management and analysis.

## Results

A total of 96,509 antibiotic prescriptions among 57,009 patient visits were retrieved from the electronic database during the study period. The ten most common diagnoses among these patients accounted for over 77% of all prescriptions, therefore we focused on only these ten diagnostic categories in the analysis.

Table 1 shows a comparison of the number of antibiotics prescribed for various common diseases stratified by appropriateness of use. Diseases of the respiratory system accounted for about 70% of all antibiotic prescriptions followed by diseases of the digestive system (13.5%) and genitourinary system (5.7%). Inappropriate use of antibiotics was found in 91.6% of the 74,648 prescriptions (7.1% for escalated use and 84.4% for incorrect and/or unnecessary use). The highest rate of inappropriate use was seen for symptoms, signs and abnormal clinical and laboratory finding not elsewhere classified (100%), diseases of the eye and adnexa (99%), and injury, poisoning and certain other consequences of external causes (98%). Escalated use of antibiotics was common for patients with disorders of the urinary system (70.0%), urethritis and urethral syndrome (68.4%) and cholecystitis (62.6%) whereas incorrect and/or unnecessary use was common to all diseases. The highest rate of appropriate use was found in diseases of pulp and periapical tissues (71.2%), acute pharyngitis (69.7%), and arthritis (67.6%).

**Table 1.** Distribution of antibiotic prescriptions stratified by clinical diagnosis and appropriateness of use

| Diagnosis | Total | Appropriate use<br>n (%) | Inappropriate use, n (%) |  |
| --- | --- | --- | --- | --- |
|  |  |  | Escalated use | Incorrect and/or unnecessary use |
| **1. Diseases of the respiratory system** | **52,186** | **4,248 (8.1)** | **1,669 (3.2)** | **46,269 (88.7)** |
| *Upper respiratory tract infection* |  |  |  |  |
| Acute tonsillitis | 4,423 | 2,193 (49.6) | 859 (19.4) | 1,371 (31.0) |
| Acute pharyngitis | 1,369 | 954 (69.7) | 0 (0) | 415 (30.3) |
| Chronic sinusitis | 332 | 81 (24.4) | 67 (20.2) | 184 (55.4) |
| Others | 34,280 | 0 (0) | 0 (0) | 34,280 (100) |
| *Lower respiratory tract infection* |  |  |  |  |
| Bronchitis, not specified as acute or chronic | 1,725 | 887 (51.4) | 333 (19.3) | 505 (29.3) |
| Pneumonia, organism unspecified | 367 | 133 (36.2) | 196 (53.4) | 38 (10.4) |
| Others | 9,476 | 0 (0) | 0 (0) | 9,476 (100) |
| **2. Diseases of the digestive system** | **10,068** | **987 (9.8)** | **2,564 (25.5)** | **6,517 (64.7)** |
| Gingivitis and periodontal diseases | 1,776 | 622 (35.0) | 138 (7.8) | 1,016 (52.2) |
| Diseases of pulp and periapical tissues | 118 | 84 (71.2) | 4 (3.4) | 30 (25.4) |
| Gastritis and duodenitis | 4,405 | 0 (0) | 973 (22.1) | 3,432 (77.9) |
| Cholecystitis | 511 | 273 (53.4) | 171 (62.6) | 67 (13.1) |
| Acute appendicitis | 49 | 8 (16.3) | 20 (40.8) | 21 (42.9) |
| Other non-infective gastroenteritis and colitis | 1,755 | 0 (0) | 985 (56.1) | 770 (43.9) |
| Others | 1,181 | 0 (0) | 0 (0) | 1,181 (100) |
| **3. Diseases of the genitourinary system** | **4,272** | **365 (8.5)** | **1,386 (32.5)** | **2,521 (59.0)** |
| Other female pelvic inflammatory disease | 1367 | 0 (0) | 567 (41.5) | 800 (58.5) |
| Other inflammation of vagina and vulva | 455 | 284 (62.4) | 0 (0) | 171 (37.6) |
| Cystitis | 166 | 0 (0) | 103 (62.1) | 63 (38.0) |
| Urethritis and urethral syndrome | 383 | 0 (0) | 262 (68.4) | 121 (31.6) |
| Other disorders of urinary system | 533 | 81 (15.2) | 373 (70.0) | 79 (14.8) |
| Others | 1287 | 0 (0) | 0 (0) | 1,287 (100) |
| **4. Diseases of the musculoskeletal system and connective tissue** | **2,318** | **669 (28.9)** | **99 (4.2)** | **1,550 (66.9)** |
| Other arthritis | 989 | 669 (67.6) | 99 (10.0) | 221 (20.9) |
| Others | 1329 | 0 (0) | 0 (0) | 1,329 (100) |
| **5. Injury, poisoning and certain other consequences of external causes** | **846** | **0 (0)** | **18 (2.1)** | **828 (97.9)** |
| Other and unspecified injuries of abdomen, lower back and pelvis | 180 | 86 (47.8) | 18 (10.0) | 76 (42.2) |
| Others | 752 | 0 (0) | 0 (0) | 752 (100) |
| **6. Diseases of the eye and adnexa** | **1,850** | **18 (1.0)** | **0 (0)** | **1,832 (99.0)** |
| Hordeolum and chalazion | 141 | 18 (12.8) | 0 (0) | 123 (87.2) |
| Others | 1647 | 0 (0) | 0 (0) | 1,647 (100) |
| **7. Diseases of the skin and subcutaneous tissue** | **949** | **25 (2.6)** | **23 (2.4)** | **901 (95.0)** |
| Cellulitis | 65 | 25 (38.5) | 23 (35.4) | 17 (26.2) |
| Others | 881 | 0 (0) | 0 (0) | 881 (100) |
| **8. Diseases of the circulatory system** | **952** | **103 (10.8)** | **0 (0)** | **849 (89.2)** |
| Rheumatic fever without mention of heart involvement | 174 | 103 (59.2) | 0 (0) | 71 (40.8) |
| Others | 778 | 0 (0) | 0 (0) | 778 (100) |
| **9. Certain infectious and parasitic diseases** | **623** | **60 (9.6)** | **112 (18.0)** | **451 (72.4)** |
| Other gastroenteritis and colitis of infectious and unspecified origin | 242 | 60 (24.4) | 112 (44.7) | 70 (28.5) |
| Others | 381 | 0 (0) | 0 (0) | 381 (100) |
| **10. Symptoms, signs and abnormal clinical and laboratory finding not elsewhere classified** | **1,091** | **0 (0)** | **0 (0)** | **1,091 (100)** |
| Abdominal and pelvic pain + fever of unknown origin + cough + pain in throat and chest + headache | 1,091 | 0 (0) | 0 (0) | 1,091 (100) |
| **Total prescriptions** | **74,648** | **6,297 (8.4)** | **5,325 (7.1)** | **63,026 (84.4)** |

Table 2 summarizes appropriateness of antibiotic use by group. Over a half (51.5%) of the prescribed antibiotics were penicillins. However, 85.3% of penicillins were incorrectly or unnecessarily prescribed. The percentage of cephalosporins and macrolides prescribed was about 14% each, of which about 84% were incorrectly or unnecessarily prescribed. Quinolones, which should be used mainly as second-line antibiotics, was found in around 6% of all prescriptions, of which 46.2% were escalated. Lincosamides, nitroimidazoles, and aminoglycosides were not commonly seen, however almost all were prescribed inappropriately.

**Table 2.**
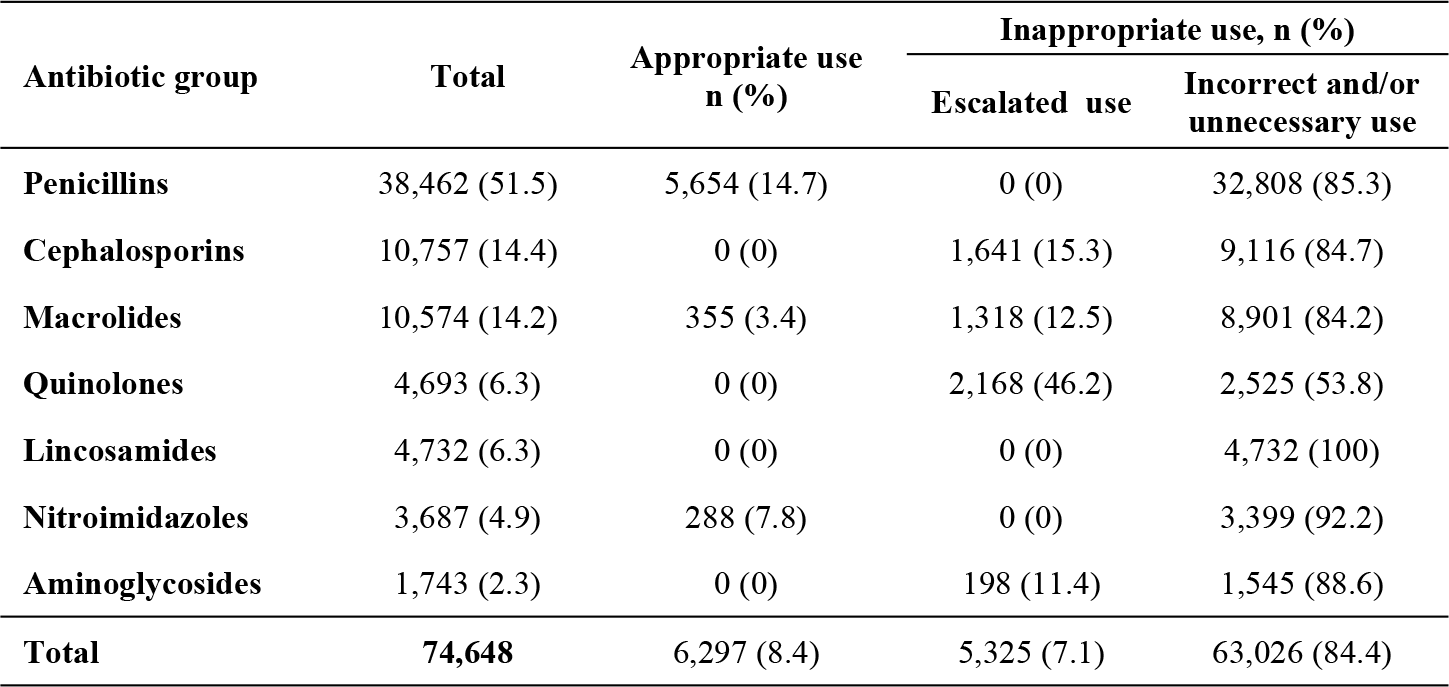
Distribution of antibiotic prescriptions stratified by antibiotic group and appropriateness of use

| Antibiotic group | Total | Appropriate use<br>n (%) | Inappropriate use, n (%) |  |
| --- | --- | --- | --- | --- |
|  |  |  | Escalated use | Incorrect and/or<br>unnecessary use |
| **Penicillins** | 38,462 (51.5) | 5,654 (14.7) | 0 (0) | 32,808 (85.3) |
| **Cephalosporins** | 10,757 (14.4) | 0 (0) | 1,641 (15.3) | 9,116 (84.7) |
| **Macrolides** | 10,574 (14.2) | 355 (3.4) | 1,318 (12.5) | 8,901 (84.2) |
| **Quinolones** | 4,693 (6.3) | 0 (0) | 2,168 (46.2) | 2,525 (53.8) |
| **Lincosamides** | 4,732 (6.3) | 0 (0) | 0 (0) | 4,732 (100) |
| **Nitroimidazoles** | 3,687 (4.9) | 288 (7.8) | 0 (0) | 3,399 (92.2) |
| **Aminoglycosides** | 1,743 (2.3) | 0 (0) | 198 (11.4) | 1,545 (88.6) |
| **Total** | **74,648** | **6,297 (8.4)** | **5,325 (7.1)** | **63,026 (84.4)** |

Table 3 compares the distribution of appropriateness of antibiotics use by physicians’ and patients’ characteristics. In the first column, “combined use” refers to physicians prescribing more than one group of antibiotic for the same patient in the same visit. The percentage of appropriate antibiotics use was reduced from that shown in Table 1 because two or more antibiotics prescribed to the same patient on the same day was considered as one visit. The percentage of each prescription type was thus appropriate use (8.7%), combined use (7.8%), escalated use (6.2%) and incorrect and/or unnecessary use (77.3%).

The odds ratios in the third column indicates the strengths of the association between inappropriate use and physicians’ and patients’ characteristics. Male physicians were more likely to prescribe antibiotics inappropriately as were those aged 32-38 years (compared to those aged less than 32 years), those who had completed a college degree, worked for 11-30 years (compared to those who had worked for less than 5 years), and were associate chief physicians (compared to resident physicians). Patients who were female, aged 18 years or more (compared to those aged ≤5 years), received antibiotics intravenously and received financial assistance from an insurance scheme were more likely to be prescribed antibiotics inappropriately.

**Table 3.**
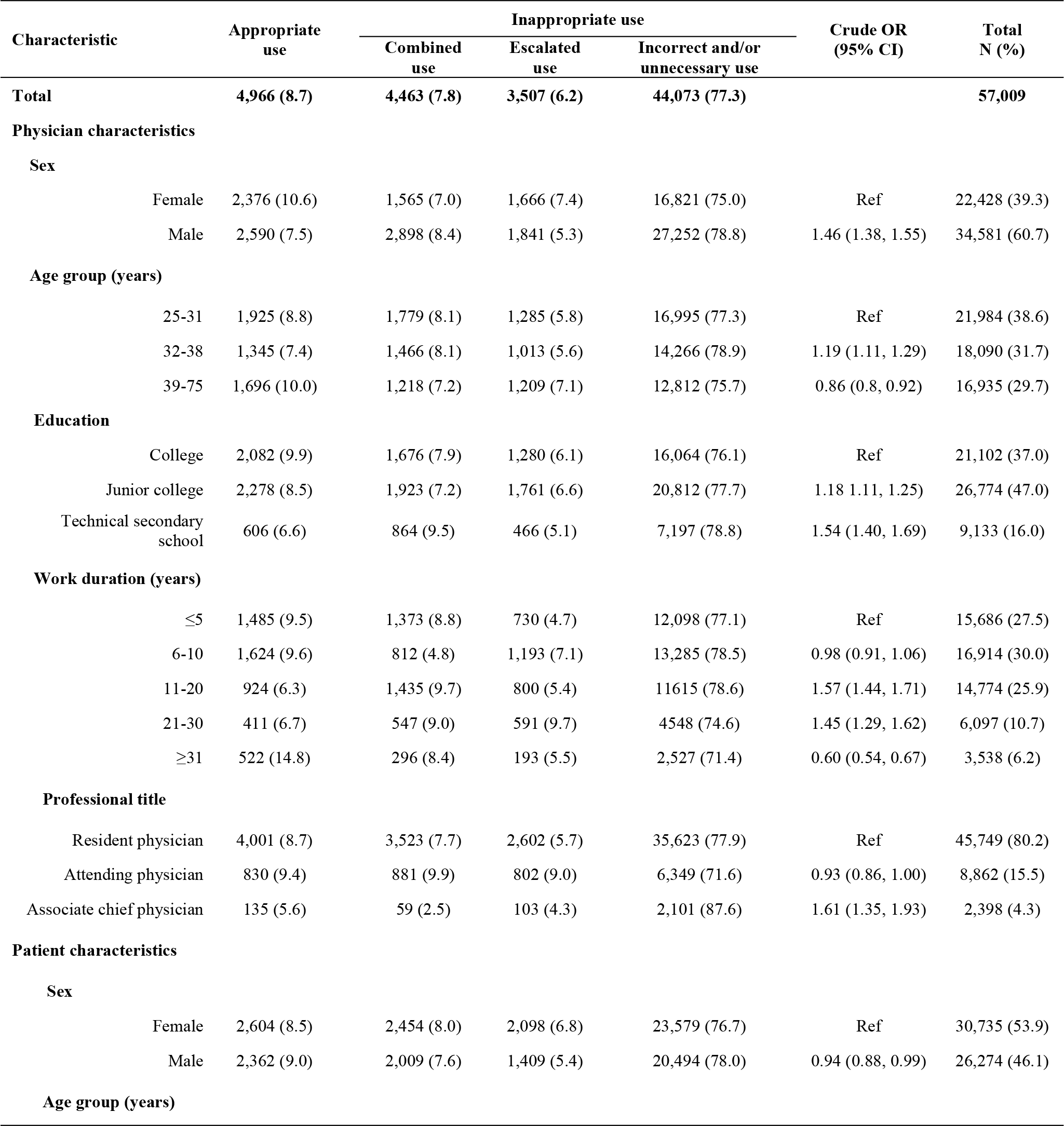
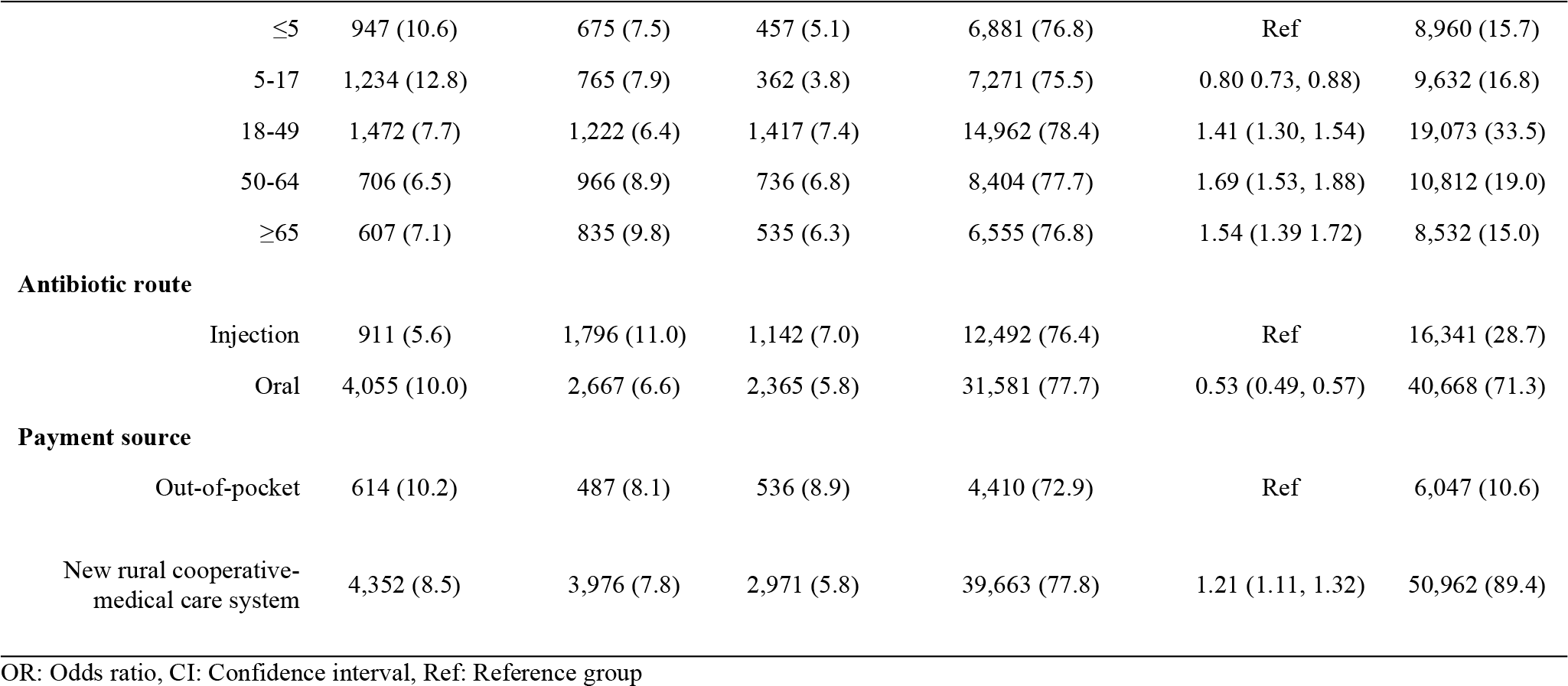
Factors associated with inappropriate use of antibiotics on univariate analysis

Table 4 shows factors associated with inappropriate antibiotic use on multivariate analysis. On the physician's side, being male, aged less than 32 years, having a lower level of education, and being an associate chief physician were associated with a higher likelihood of inappropriate antibiotic use. Increasing work duration did not equate to a higher odds of inappropriate antibiotic use. However, compared to newly employed physicians, those who had a work experience duration of more than 5 years were more likely to prescribe antibiotics inappropriately. Antibiotics prescribed intravenously were more likely to be inappropriate. On the patient's side, younger patients (age≤17 years) were more likely to get appropriate prescription. Finally, patients who received financial assistance from an insurance scheme were more likely to be prescribed antibiotics inappropriately.

**Table 4.**
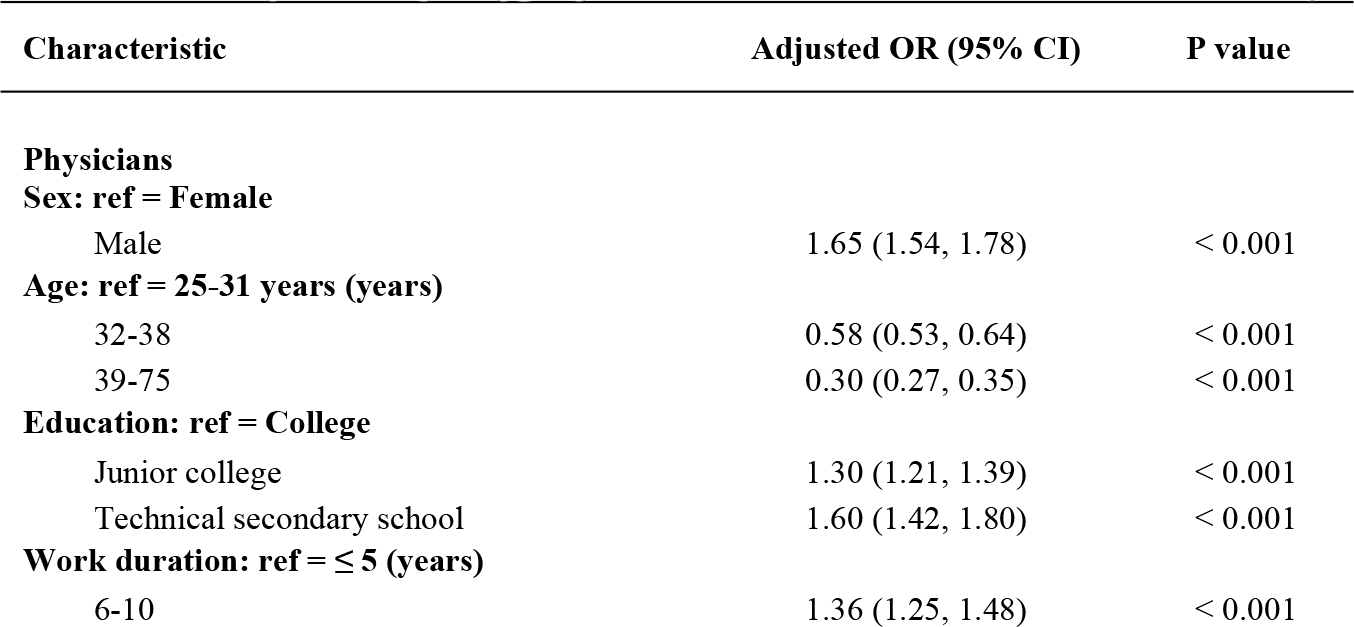
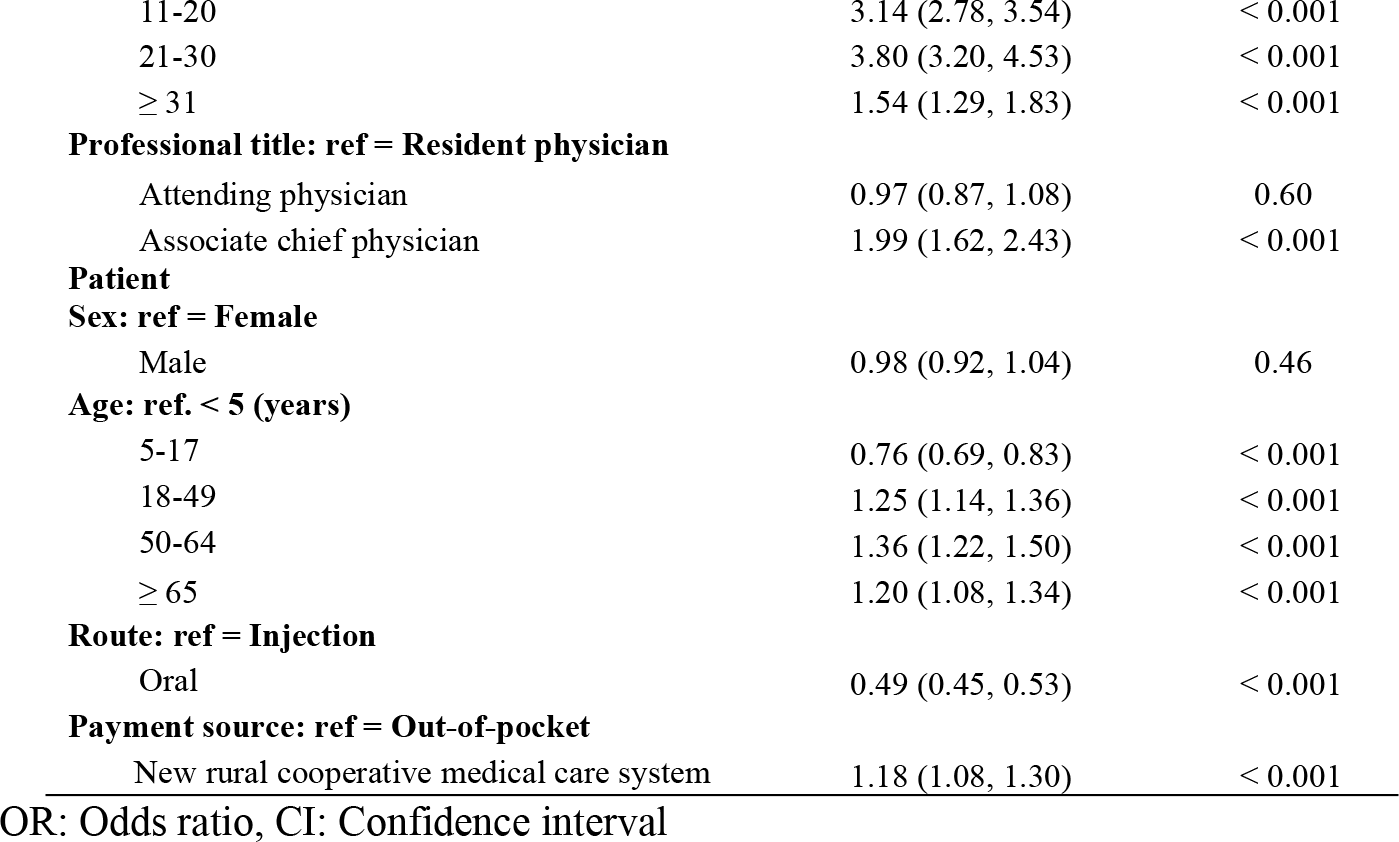
Factors predicting inappropriate use of antibiotics on multivariate analysis

## Discussion

This study showed four patterns of antibiotic use, namely: 1) appropriate use, 2) incorrect and/or unnecessary use, 3) escalated use, and 4) combined inappropriate use. More than 90% of antibiotic prescriptions were inappropriate and incorrect and/or unnecessary use was the most common pattern. Respiratory infection was the most common disease linked with antibiotic prescriptions. While penicillins were the most commonly prescribed antibiotic, cephalosporins and macrolides were more inappropriately prescribed, particularly due to escalation. These antibiotics could have been de-escalated to penicillins. The background education of these primary care physicians was generally below college level. A physician's low level of education and senior position were significantly associated with inappropriate antibiotic prescription. Antibiotic misuse was also associated with injection route as well as adult patients and those who had insurance.

In general, uncomplicated respiratory infections are mostly caused by viruses, which cannot be treated with antibiotics [13]. Unnecessary use of antibiotics to treat such infections has been reported in many countries such as Sri Lanka, USA, Qatari, Korea Canada and China [15–20]. In our study, this accounted for more than a half of the prescriptions. Overuse of antibiotics could increase the duration of the disease and associated costs and increase resistance to infections as well as increase the risk of adverse drug reactions [21].

Penicillins are recommended for skin infections since the disease is mostly caused by penicillin sensitive bacteria [22]. Cephalosporins have a broad spectrum effect and therefore should be reserved for empirical use in patients with serious undifferentiated infections. Fluoroquinolones as well as macrolides are prescribed for those with respiratory tract infections with a history of penicillin allergy. Aminoglycosides are inappropriate for respiratory tract infections due to their toxicity as well as being more suitable for gram negative bacterial infections. In our study, although penicillins were the most commonly used antibiotic, there were cases where penicillins should have been used but were inappropriately replaced by other antibiotics.

As much as 29% of antibiotics used in this study was administered by injection. Overuse of injections was also seen other countries such as Vietnam, India, and Korea [19, 23, 24]. This practice is rarely needed in primary care as most infections can be controlled with oral antibiotics [25]. Injection of drugs is complicated by serious adverse drug reaction and diseases complications such as local infections, bleeding, and nerve injury [19, 26]. There are many guidelines and punitive measures in China that guide physicians on how to use antibiotics, but there are no such guidelines as how physicians should talk to patients and families about whether antibiotics are needed or not, and discuss possible harms. Furthermore, there are no measures to teach patients how to use antibiotics and manage symptoms of non-bacterial infections [21].

Most of the antibiotic prescriptions in this study were made by resident physicians with a below college level of education, and this was associated with antibiotic overuse and misuse. Based on this evidence, refresher courses on antibiotic prescribing for primary care physicians are necessary [7, 9]. The training should emphasize avoidance of incorrect and unnecessary use, narrow-spectrum antibiotics use, and when to prescribe injection antibiotics.

The whole analysis of this study was based on data keyed in by the primary physicians. While the antibiotic information is very accurate, that on disease classification might be less so. Accurate diagnosis is a prerequisite of appropriate selection of antibiotic. Data on appropriateness of antibiotic choice in this study must therefore be interpreted with caution.

## Conclusion

Overuse of antibiotics for uncomplicated respiratory infections, use of cephalosporins, macrolides and injection antibiotics in primary care are the major problems of clinical practice in the study areas.

## Acknowledgments

The authors thank all members of the investigational team who collected the data. We also thank all of the participating institutions for providing information and assistance during the study.

## Funding

This study is part of the first author’s thesis in partial fulfillment of the requirements for a PhD at Prince of Songkla University, Thailand, and the China Medical Board under the project “A second collaborative program to improve the health research capacity of western medical universities in China and Prince of Songkla University (PSU)”. This study was also possible due to a grant obtained from Guizhou innovative talent foundation (2016-4015).

## Disclosure

The author reports no conflicts of interest in this work.

